# Reduced signatures of gene duplication and non-random gene organization in shaping stage-specific patterns of gene expression across a relatively simple life cycle

**DOI:** 10.1101/2024.12.21.629888

**Authors:** James G. DuBose, Levi T. Morran

## Abstract

Complex eukaryotes generally experience some degree of ecological transition as they develop, which is usually accompanied by the expression of different traits across their ontogeny. Theory suggests the evolution of differentiation between life stages is facilitated by the expression of different genes at different points throughout ontogeny, which alleviates pleiotropic evolutionary constraints. Therefore, ascertaining what contributes to specialized patterns of gene expression across ontogeny is fundamental to understanding the evolution of ontogenetic complexity. Expression divergence between duplicate genes and gene organization on chromosomes have been identified as important features of relatively complex ontogenies. Therefore, one could predict that their link to transcriptional specialization across ontogeny may be weaker in relatively simpler life cycles. Here, we investigated the links between gene duplication, gene organization, and stage-specific patterns of expression across the relatively simple *Caenorhabditis elegans* life cycle. We found that signatures of stage-biased chromosomal regions were weaker in *C. elegans* than what has been previously described in organisms with more complex ontogenies. Furthermore, we found the extent that duplicate genes varied in ontogenetic expression pattern was more constrained in *C. elegans*. Overall, our findings add to a body of evidence that suggests a link between gene duplication and gene organization and the evolution of complex ontogenies.

## Introduction

Complex eukaryotes generally experience some degree of ecological transition as they develop, which is usually accompanied by the expression of different traits across their ontogeny. Prominent questions in evolutionary biology often pertain to how individual genomes evolve to encode these varieties of traits. Broad interest in these questions has fueled developmental evolutionary biology, which has yielded general predictions of genome evolution in light of ontogenetic dynamics. Theory suggests that pleiotropy, where the same gene produces traits in different ecological and developmental contexts, should constrain the diversification of phenotypes encoded in the same genome (Williams 1957; Ebenman 1992; Moczek et al. 2011). For example, the probability that a given genetic variation is adaptive or neutral is lower when expressed in multiple life stages than when expressed in a single life stage because there are more chances for it to have deleterious effects (Ebenman 1992). However, this issue can be circumvented through the expression of different genes in different ontogenetic contexts, which can confer a greater degree of evolutionary independence between life stages (Haldane 1932; Ebenman 1992). A growing body of theory and empirical studies is clarifying our understanding of the interplay between ontogenetic patterns of gene expression and evolutionary processes. However, previous studies have predominately focused on organisms with relatively complex ontogenies and questions remain of how generalizable their findings are to organisms with relatively less complex ontogenetic changes.

Understanding what coordinates and changes patterns of gene expression across ontogeny is fundamental to understanding the evolution of differentiation between life stages. Expression divergence between duplicate genes is thought to be an integral part of this process (Li et al. 2005; DuBose and de Roode 2024; Ma et al. 2024). Gene duplication provides a rich source of genetic material to facilitate significant evolutionary changes (Ohno, Susumu 1970; Zhang 2003). Furthermore, gene duplication can alleviate pleiotropic constraints by allowing for greater specialization in patterns of expression (Li et al. 2005; Ma and Zheng 2023; Ma et al. 2024). However, patterns of gene expression are often coordinated by more complex regulatory features that operate independent of specific gene sequences. In eukaryotes, co-expression of non-randomly localized genes on chromosomes is among the most prominent of these features (Hurst et al. 2004; Zinani et al. 2022). Previous studies have predominately focused on co-expression of localized genes as it pertains to tissue-specificity and gene function (Boutanaev et al. 2002; Lercher et al. 2002; Williams and Bowles 2004). However, the role of gene localization in promoting modular expression patterns between life stages across ontogeny is less understood.

Recent evidence suggests that localization of genes with shared stage-biased expression patterns may be a general feature of holometabolous development (complete metamorphosis) in insects (Wojciechowski et al. 2018; Kimura et al. 2024). Holometabolous development involves significant morphological and physiological restructuring, placing it among the most complex ontogenetic programs. However, the roles of expression divergence between duplicate genes and gene organization in shaping relatively simpler ontogenies are less understood. If expression specialization of duplicate genes and gene organization are important features for the evolution of more complex ontogenetic programs, one could predict that their link to transcriptional specialization across ontogeny may be weaker in relatively simpler life cycles.

To begin exploring this idea, we examined the links between gene organization, gene duplication, and stage-specific patterns of expression across the relatively simple *Caenorhabditis elegans* life cycle. The *C. elegans* life cycle progresses through four larval stages (L1-L4) before reaching adulthood. When developing in stressful conditions, *C. elegans* individuals express dauer larvae, which is a plastic alternative third larval stage that is more robust to stressful conditions (Riddle et al. 1981). The interplay between gene duplication and the evolution of *C. elegans* ontogenetic dynamics has been well studied (Castillo-Davis and Hartl 2002; Konrad et al. 2018; Lu et al. 2020; Ma and Zheng 2023; Ma et al. 2024). Here, we expand on this previous work by comparing the ontogenetic expression dynamics of duplicate genes across the *C. elegans* life cycle to more complex holometabolous insect ontogenies. A role of non-random gene organization in coordinating patterns of *C. elegans* gene expression has been described as well (Kim et al. 2001; Lercher et al. 2003; Cutter et al. 2009). However, this has predominately been done in the contexts of embryonic development, tissues-specificity, and gene function similarities. Therefore, here we expand on these previous studies by quantifying non-random arrangements of stage-biased genes on chromosomes across the *C. elegans* life cycle.

## Methods

### Data accession and processing

To quantify *C. elegans* ontogenetic expression patterns, we used existing time-series gene expression data from (Boeck et al. 2016), which normalized expression values using depth of coverage per base per million reads normalization. We then averaged expression values and log transformed them prior to downstream analyses. To compare the expression dynamics of duplicate genes between *C. elegans* and the more complex holometabolous insect ontogeny, we used available time-series gene expression data from the fruit fly *Drosophila melanogaster* and the monarch butterfly *Danaus plexippus*. We used *D. melanogaster* temporal gene expression data from FlyBase (Graveley et al. 2011; Öztürk-Çolak et al. 2024) and *D. plexippus* temporal gene expression data from (DuBose and de Roode 2024). These data were normalized using transcript per million normalization, and we subsequently processed them the same way as the *C. elegans* data. We note that the difference in normalization procedure is unlikely to influence our inferences because all of our comparisons were performed using the τ index (described below), which calculates expression bias of a particular gene relative to its expression in life stages. In other words, quantification of τ only involves within dataset comparison, not between datasets.

### Algorithm for identifying stage-biased regions

Before identifying stage-biased regions, we first calculated each gene’s level of stage-specificity using the τ index (Yanai et al. 2005). τ ranges from 0 (even expression across life stages) to 1 (expression is isolated to single life stage): , where *N* represents the number of life stages and *xi* is the expression level normalized to the maximum expression across stages. To search for stage-biased regions, we used a sliding-window algorithm that identifies stage-biased regions by quantifying the standardized difference (Cohen’s d) between the stage-specificity of genes within a window to a random sample of 1000 genes from across the corresponding chromosome. A Cohen’s d value of greater than or equal to 1 indicates that genes within a given window were significantly more stage-specific than a representative sample from the rest of their chromosome, which was our criterion for defining stage-biased regions. We used this algorithm on each life stage and used sliding window sizes of 100kb and 10kb (which did not impact our general inferences, see supplemental section 1.2).

### Testing for non-randomness of the number of and gene density in stage-biased regions

After identifying stage-biased regions, we were then interested in if the number and average gene density of these regions was greater than expected if expression patterns were randomly distributed among genes. First, we derived expected null distributions for the number of stage-biased regions and their mean gene density using Monte-Carlo simulations that randomly shuffled expression patterns among genes and used the previously described algorithm to identify stage-biased regions. We ran the simulation for 1000 iterations, and for each iteration, we calculated the number of stage-biased regions and their mean gene density for each life stage. We then calculated the proportion of each null distribution that was greater than or equal to the observed values (which we report as P(N≥O)).

After identifying stage-biased regions, we were interested in their contribution to general patterns of transcriptional specification between stage. We quantified the proportion of stage-biased genes that were in stage-biased regions as a function of their stage-specificity (τ) by fitting logistic regression models. Note that no L3-biased regions were detected, and there was not a sufficient number of genes in L2-biased regions for fitting a logistic model. We fit logistic models using the g*lm* R function with a binomial link function (R Core Team 2022) (see Table S1 for full model summaries).

### Examining patterns of gene duplication in stage-biased regions

Since patterns of expression may be conserved in duplicate genes, we quantified the extent that stage-biased regions were comprised of duplicate genes. To identify homologous gene groups, we first used PSI-BLAST (BLAST 2.5.0+) (Altschul 1997) with 5 iterations to align *C. elegans* protein sequences to each other (WormBase release WS294, GenBank Assembly GCA_000002985.3). We then inferred genes to be homologous if the query sequence showed at least 30% similarity across the length of the target sequence and an E-value of at least 1 x 10^-10^, which was also the set of criteria previously used for identifying the *D. plexippus* homologous groups we used for comparisons (DuBose and de Roode 2024). Finally, we grouped homologous pairs into two-node subgraphs and merged subgraphs based on common node identity to fully assemble homologous gene groups.

After identifying groups of homologous genes, we calculated the number of unique homologous groups represented in each stage-biased regions divided by its gene density, which we refer to as the non-homologous ratio. Here, a value of 1 indicates that a given stage-biased region is comprised entirely of non-homologous genes, and lower values indicate that the stage-biased region is largely comprised of duplicates. We then conducted a correlation test between non-homologous ratio and the density of genes within each stage-biased region using the Spearman rank correlation. We used the *cor.test* R function for statistical testing (R Core Team 2022), and visualized the non-parametric correlation by fitting a local regression using the *lowess* function from the *statsmodels* Python library (Seabold and Perktold 2010).

### Comparing divergence in patterns of stage-specificity within homologous groups between C. elegans and the holometabolous insects

To gain further insight into the role of gene duplication in facilitating transcriptional differentiation between life stages, we compared the degree of conservation in patterns of stage-bias and stage-specificity (τ) within *C. elegans* homologous groups to the holometabolous insects *D. melanogaster* and *D. plexippus*. We first identified homologous groups for *D. melanogaster* (GenBank assembly GCA_000001215.4) and *D. plexippus* (GenBank assembly GCA_000235995.2) as previously described for *C. elegans*. Next, we quantified the standard deviation in stage-specificity within each homologous group. We then assigned a stage-bias to each gene in each homologous group based on the stage of their maximum expression and calculated the proportion of the homologous group that shared the most frequent stage-bias. This metric is equivalent to the standard Berger-Parker dominance index frequently used in community ecology, but we refer to this as stage-bias uniformity for our purposes. A stage-bias uniformity value of 1 indicates that all genes within a homologous groups are biased towards the same stage, and lower values indicate that many different stage-biases are represented in the homologous group. We tested for differences in the distributions of τ standard deviation and stage-bias uniformity between *C. elegans* and both *D. melanogaster* and *D. plexippus* homologous groups with Kolmogorov-Smirnov tests using the *ks_2samp* function from the *scipy.stats* Python library (Virtanen et al. 2020).

## Results

### Signatures of non-random stage-biased chromosomal regions vary by life stage

We detected stage-biased regions for each *C. elegans* life stage with the exception of L3 (Figure 1A, Figure S1). The number of these stage-biased was significantly greater than expected if expression patterns were randomly distributed for the embryo, L1, dauer, and adult stages (P(N≥O) ≤ 0.047) (Figure 1B). However, the number of stage-biased regions were consistent with null predictions for L2, L3, and L4 (P(N≥O) ≥ 0.051) (Figure 1B). We also found that the mean density of genes in stage-biased regions was significantly greater than expected for the embryo, L1, dauer, and L4 stages (P(N≥O) ≤ 0.047) (Figure 1C). However, the mean density of genes in stage-biased regions was consistent with null predictions for the L2, L3, and adult stages (P(N≥O) ≥ 0.058) (Figure 1C). We then fit logistic models to examine the proportion of genes in stage-biased regions as a function of their stage-specificity. We found that although the proportion of genes found in stage-biased regions increased with their stage-specificity (log-odds ≥ 5.3, p ≤ 1.14 x 10^-4^) , the overall proportion varied by life stage and was relatively low for any given class of stage-specificity (Figure 2).

**Figure 1.**
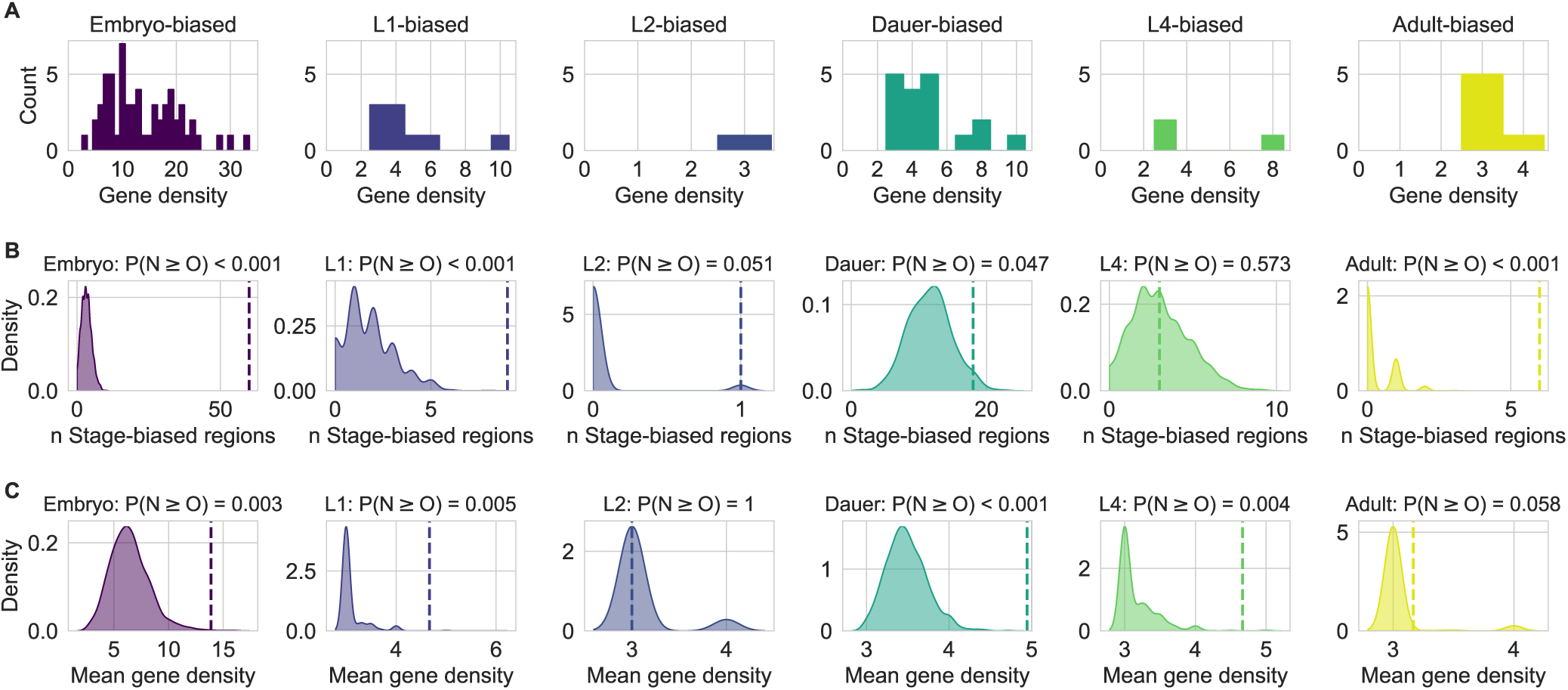
Signatures of non-random stage-biased chromosomal regions vary by life stage. A) Each histogram represents the number of stage-biased regions as a function of the number of stage-biased genes in the region (gene density). B and C) The shaded region in each panel shows the expected distributions represented as kernel density estimates of B) the number of stage-biased regions, and C) the mean gene density within stage-biased regions if expression patterns were randomly distributed. Vertical dashed lines indicate the observed values, and the text above each panel indicates the probability that the observed value was produced by the null simulations.

**Figure 2.**
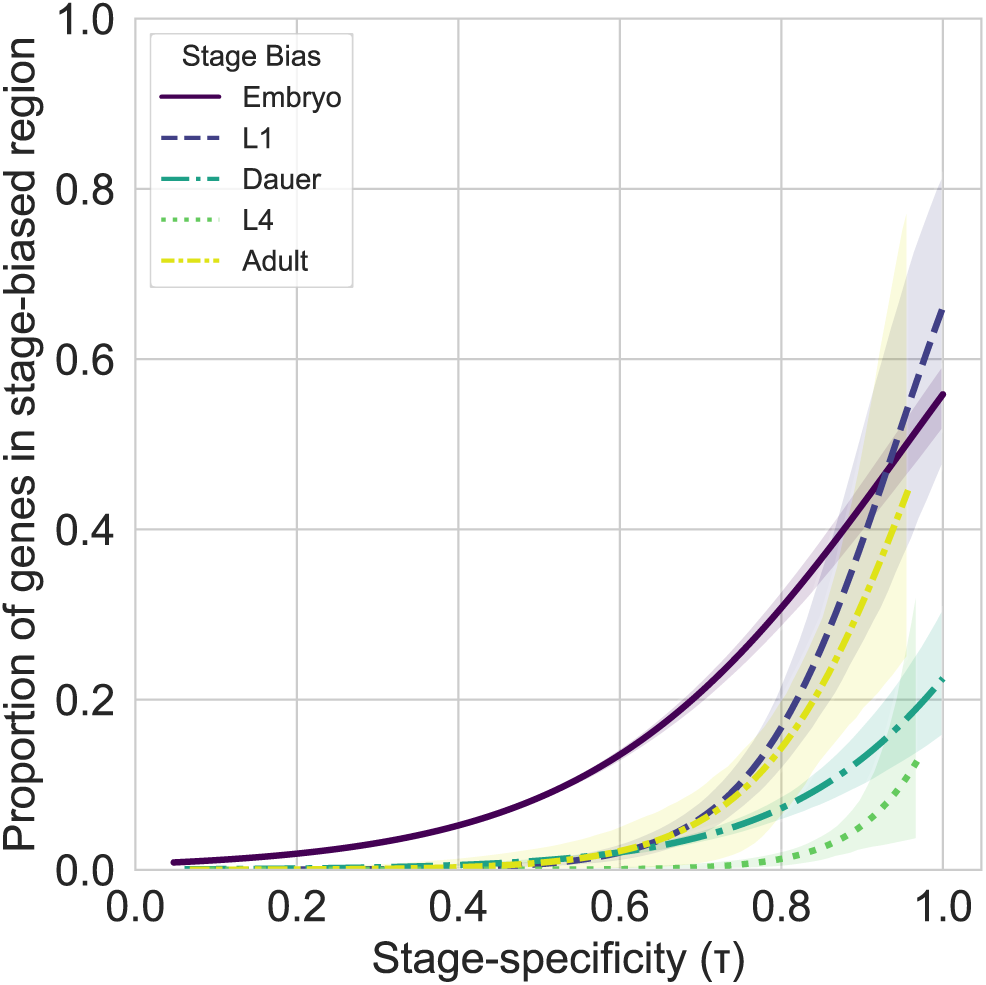
The proportion of stage-specific genes that are in stage-biased increases with stage-specificity and varies between life stages. Each line represents the logistic model that describes the proportion of genes that are in stage-biased regions (y-axis) as a function of stage-specificity (x-axis). Shaded regions around the lines represent the 95% confidence interval.

### The contribution of gene duplication to the formation of stage-biased regions

Since local gene duplication can contribute to the formation of stage-biased regions, we quantified the proportion of homologous groups represented in each stage-biased regions relative to the number of genes. This showed that approximately 30% of stage-biased regions did not contain detected homologs (Figure 3A). The remainder of stage-biased regions contained at least one set of duplicates, but the extent of duplication varied significantly and was generally low (Figure 3A) For example, approximately half of all stage-biased regions were comprised of greater than 80% non-homologous genes (Figure 3A). We then examined the correlation between the non-homologous ratio and gene density and found that more dense stage-biased regions tended to be composed of a greater proportion of duplicate genes (ρ = -0.32, p = 0.0012) (Figure 3B).

**Figure 3.**
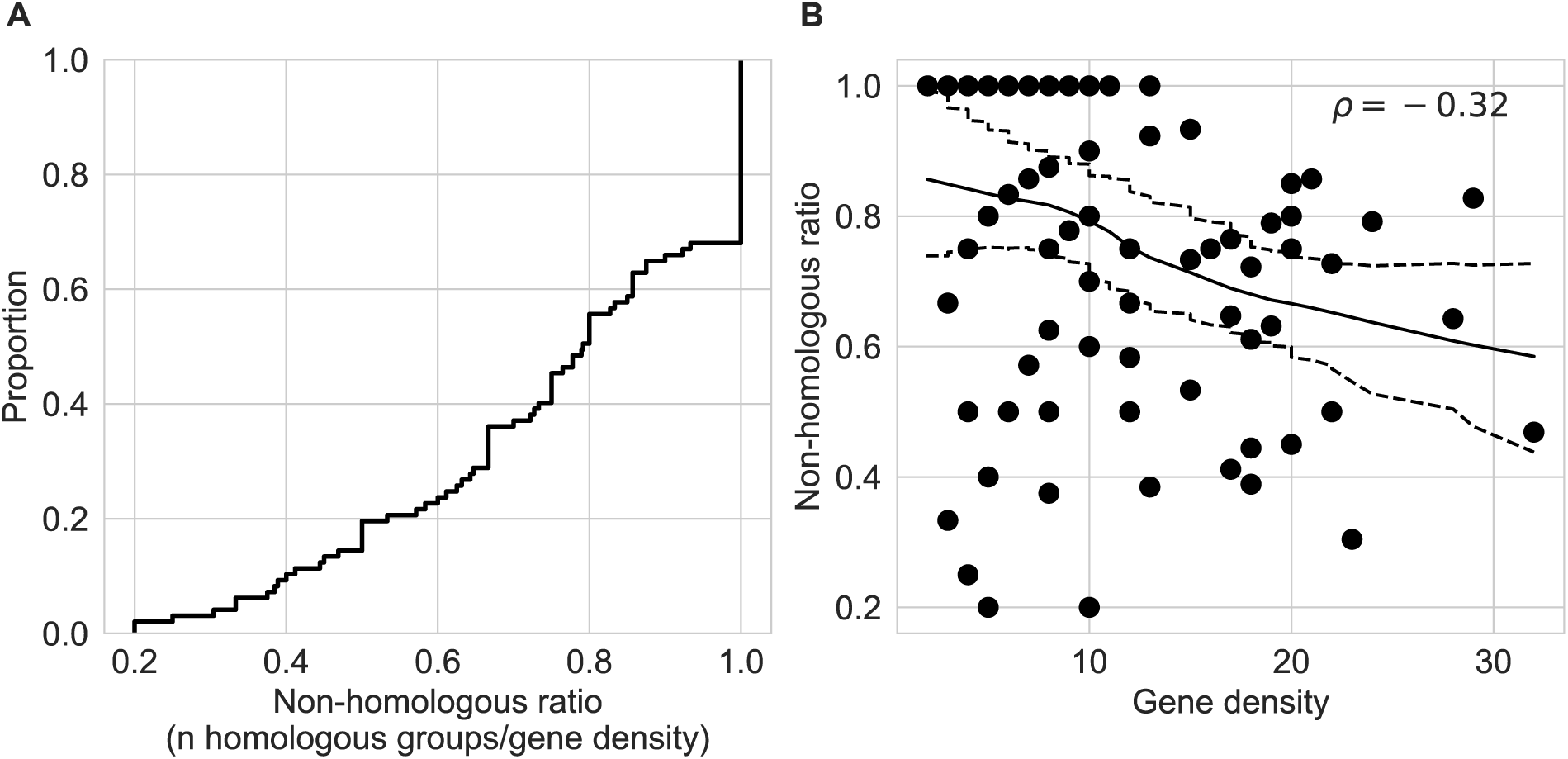
Patterns of expression similarity between genes in stage-biased regions can be partially but not entirely explained by gene duplication. A) An empirical cumulative density plot showing the distribution of non-homologous ratios across stage-biased regions. Non-homologous ratios represent the number of homologous groups represented in a stage-biased region relative to the number of genes. A non-homologous ratio of 1 indicates that all genes within a stage-biased region are non-homologous. B) Non-homologous ratios as a function of gene density for stage-biased regions. The solid and dashed lines represent the fit local regression model and 95% confidence interval, respectively. Note that the fit model is used to visualize the negative correlation, not to make statistical inferences.

### Change in ontogenetic expression pattern between duplicate genes is relatively constrained in C. elegans

Theory suggests that gene duplication can alleviate pleiotropic constraint. However, it must also be accompanied by changes in ontogenetic expression patterns to facilitate differentiation between life stages. To contextualize the extent that ontogenetic expression patterns change in duplicate genes, we compared the variation in expression pattern within *C. elegans* homologous groups to those of the holometabolous insects *D. melanogaster* and *D.* plexippus. We found that *C. elegans* homologous groups had a decrease standard deviation in stage-specificity (τ) relative to *D. melanogaster* (D = 0.20, p = 1.73 x 10^-5^) and *D. plexippus* (D = 0.31, p = 2.39 x 10^-10^) (Figure 4A). We then calculated the proportion of genes within a given homologous group that shared the most frequent stage-bias. This showed that expression patterns within *C. elegans* homologous groups were more often biased towards a single stage than those of *D. melanogaster* (D = 0.35, p = 8.2 x 10^-15^) and *D. plexippus* (D = 0.35, p = 1.3 x 10^-13^) (Figure 4B). Taken together, these findings suggest that changes in ontogenetic expression pattern between duplicate genes is relatively more constrained in *C. elegans* than in holometabolous insects.

**Figure 4.**
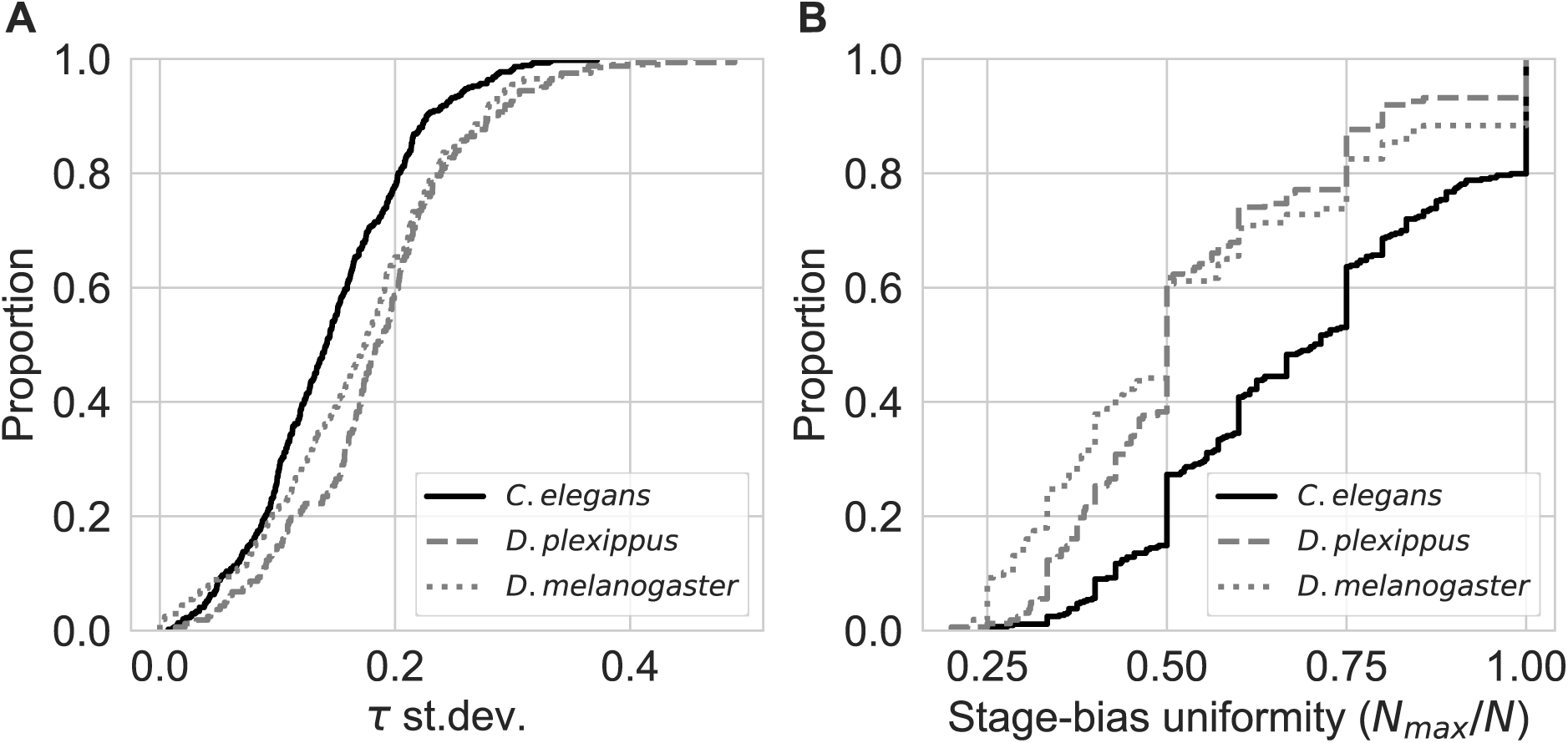
Expression pattern divergence within homologous groups is relatively constrained in *C. elegans*. A) Empirical cumulative density functions showing the distributions of variation in stage-specificity (tau) within homologous groups. The solid line represents *C. elegans,* the dashed line represents the monarch butterfly *D. plexippus*, and the dotted line represents *D. melanogaster*. There is generally less variation in stage-specificity between genes within *C. elegans* homologous groups relative to those of *D. plexippus* and *D. melanogaster*. B) Empirical cumulative density functions showing the distributions of stage-biased uniformity in *C. elegans, D. plexippus*, and *D. melanogaster* homologous groups. Stage-bias uniformity represents the proportion of genes within a homologous group that share the most frequent stage-bias, such that a value of 1 indicates that all genes share the same stage-bias.

## Discussion

Expression specialization of duplicate genes and localization of stage-biased genes on chromosomes have been identified as important features of complex ontogenetic programs. To complement existing studies, here we examined the link between these features and stage-specific patterns of gene expression across the simpler *C. elegans* life cycles. In summary, we found that the number and size (gene density) of stage-biased chromosomal regions varied between *C. elegans* life stages and were most apparent in embryo and dauer larva (Figure 1). However, the proportion of genes present in stage-biased regions was relatively low. This was also the case for even the most stage-specific genes (τ > 0.75), indicating that gene localization on chromosomes is not a general driver of transcriptional differentiation between *C. elegans* life stages (Figure 2). We then investigated the potential for gene duplication to explain the presence of the observed stage-biased regions and found that nearly 70% of stage-biased regions contained at least one set of duplicate genes (Figure 3A). However, the abundance of duplicate genes in stage biased regions was generally low and was greater in more gene dense stage-biased regions (Figure 3). This suggests that although gene duplication could play a role in the expansion of stage-biased regions, it alone does not explain their presence. Finally, we found the extent that duplicate genes vary in ontogenetic expression pattern was significantly constrained in *C. elegans* relative to the holometabolous insects *D. melanogaster* and *D. plexippus* (Figure 4).

A growing body of evidence suggests that the evolution of complex ontogenies may have been facilitated by features such as gene duplication and gene organization, which can play important roles in alleviating pleiotropic constraints (Wojciechowski et al. 2018; DuBose and de Roode 2024; Kimura et al. 2024). Our findings are consistent with this evidence but highlight the contrary. The signatures of non-random stage-biased regions that we identified in *C. elegans* are notably weaker than what has been previously identified in organisms with more complex ontogenies, such as holometabolous insects (Duncan et al. 2020; Kimura et al. 2024). This pattern is also consistent with previous findings that signatures of large-scale chromosomal features that coordinate expression, such as topologically associated domains, are relatively weak in *C. elegans* (Crane et al. 2015). Furthermore, we found that changes in ontogenetic expression pattern between duplicate genes, which has been linked to the differentiation between stages across complex life cycles, was relatively constrained in *C. elegans* (DuBose and de Roode 2024). Taken together, these findings suggest weaker roles of expression specialization between duplicate genes and gene organization in generating transcriptional differentiation between life stages across simpler ontogenies.

The life stage with the most abundant and gene dense biased regions was embryo, which is unsurprising given the significant gene expression coordination required for embryonic development (Hanchuan Peng et al. 2006; Cohen-Fix and Askjaer 2017; Jash and Csankovszki 2024). A more interesting finding was that the stage with the second most abundant and dense stage-biased regions was dauer larva. Dauer larva is a functionally unique and important life stage for *C. elegans*, and its evolution represents a significant life cycle innovation for Rhabditida nematodes (Walter Sudhaus and David Fitch 2001; Félix and Braendle 2010). Therefore, the enrichment of dauer-biased regions relative to other life stages is consistent with the notion that non-random gene organization could be associated with the evolution of greater life cycle complexity. However, the proportion of dauer-biased genes localized to dauer-biased regions was relatively low, suggesting additional factors were at play during the evolution of the dauer larva stage.

Although our findings were consistent with theory and previous evidence, we interpret them with caution. Many more studies are needed to establish the link between the evolution of ontogenetic complexity and the genetic architectural features we explored here. Furthermore, the majority of studies (including this one) that investigate the evolution of ontogenetic complexity generally focus on the genetic and transcriptional features of complex life cycles. While these studies have provided a great deal of insight into how pleiotropic constraints could be alleviated, they are inherently associative. Experimental evaluation of the interplay between pleiotropic constraint and life cycle evolution is limited and has primarily focused on organisms with relatively complex ontogenies (Fellous and Lazzaro 2011; Aguirre et al. 2014; Collet and Fellous 2019; Critchlow et al. 2019; Goedert and Calsbeek 2019). Therefore, future studies on organisms with simpler ontogenies could contribute to a more comprehensive understanding of what facilitates and constrains the evolution of ontogenetic complexity.

## Data and code availability

No original data was generated for the work presented in this manuscript. All code written for analyses has been deposited in the GitHub repository https://github.com/gabe-dubose/celga and has been archived under the DOI 10.5281/zenodo.14241740.

## Appendix for: Reduced signatures of gene duplication and non-random gene organization in shaping stage-specific patterns of gene expression across a relatively simple life cycle

### 1 Supporting results

#### 1.1 A graphical overview of stage-biased regions

**Figure S1:**
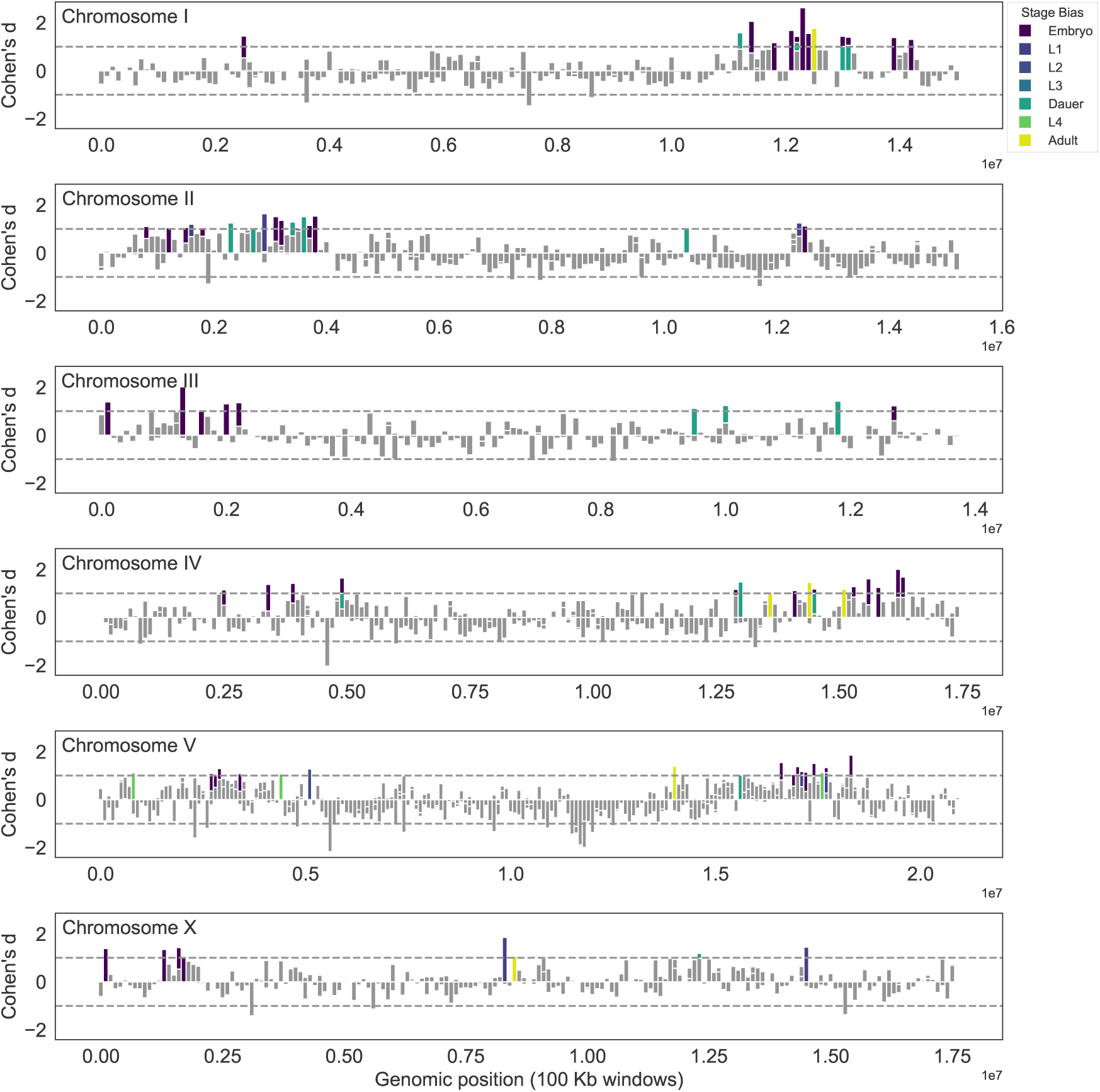
A graphical overview of stage-biased regions identified across the *C. elegans* genome. Each panel represents a chromosome. In each panel, bars represent a 100kb window and the height represents the Cohen’s d value for the corresponding window. Bars are colored by the stage in which they are biased if the Cohen’s d is *≥* 1, which indicates significant stage-bias.

#### 1.2 Stag-biased regions using a 10kb sliding window

To ensure our detection of stage-biased regions was robust to different scales, we searched for stage-biased regions using a 10kb sliding window, rather than the 100kb sliding window presented in the main text. Using this smaller sliding window, we detected stage-biased regions for all life stages besides L2 and L3, which is consistent with the analysis presented in the main text (Figure S2).

**Figure S2:**
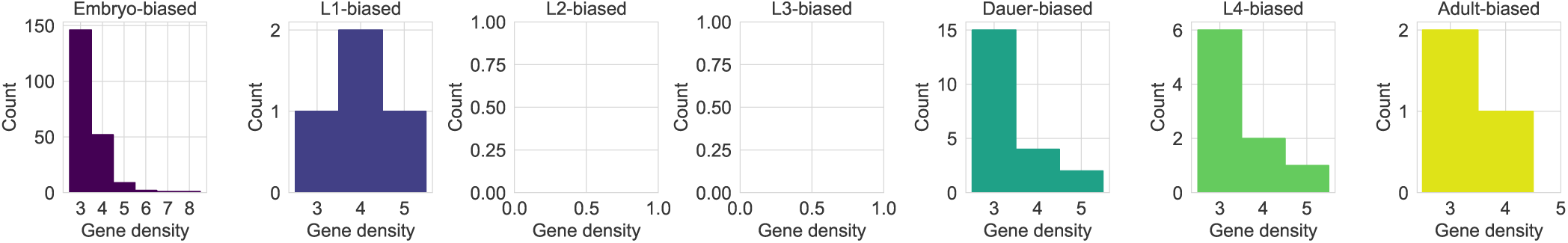
Signatures of stage-biased chromosomal regions vary by life stage. Each histogram represents the number of stage-biased regions as a function of the number of stage-biased genes in the region (gene density)

We then tested for non-randomness in the number and size (gene density) of these stage-biased regions as described in the main text. This showed that the number of these stage-biased was significantly greater than expected if expression patterns were randomly distributed for the embryo, L1, dauer, L4, and adult stages, but not for L2 and L3 (Figure S3). Here, the only difference from the main text is that the number of L4-baised regions was detected as non-random. The gene density within stage-biased regions was also significantly greater than the null expectation for embryo, L1, dauer, L4, and adult, which is consistent with the findings presented in the main text (Figure S4).

**Figure S3:**
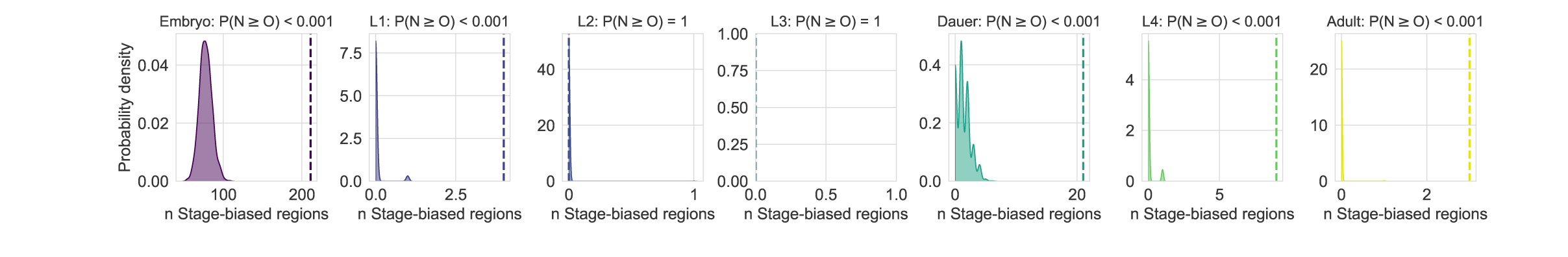
Non-randomness in the number of stage-biased regions varies by life stage. The shaded region in each panel shows the expected distribution of the number of stage-biased regions if expression patterns were randomly distributed. Panels without distributions indicate no stage-biased regions were predicted. Vertical dashed lines indicate the observed values.

**Figure S4:**
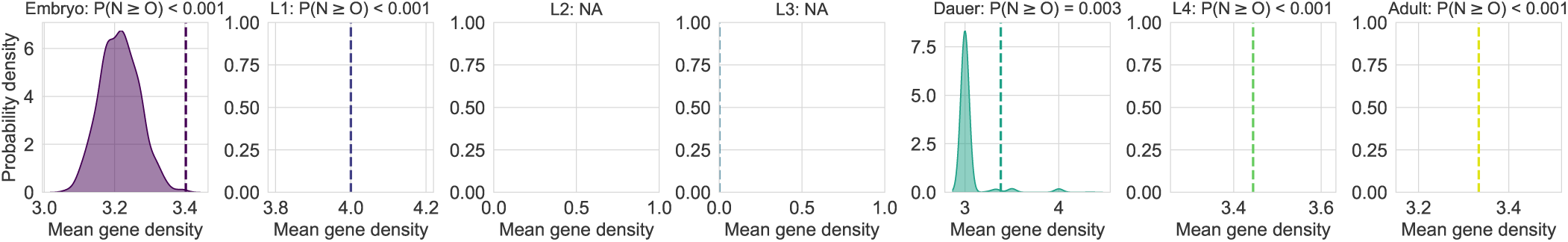
Non-randomness in the gene density of stage-biased regions varies by life stage. The shaded region in each panel shows the expected distribution of the number of stage-biased regions if expression patterns were randomly distributed. Panels without distributions indicate no stage-biased regions of sufficient density (n genes *≥* 3) were predicted. Vertical dashed lines indicate the observed values.

We then quantified the proportion of genes present in stage-biased regions as a function of stage-specificity using logistic regression models. This showed a slightly greater proportion of genes in stage-biased regions than the primary analysis (Figure S5, Table S2). However, the proportions were still generally less than half even for even the most stage-specific genes (with the exception of embryo). This analysis is largely consistent with what is presented in the main text.

**Figure S5:**
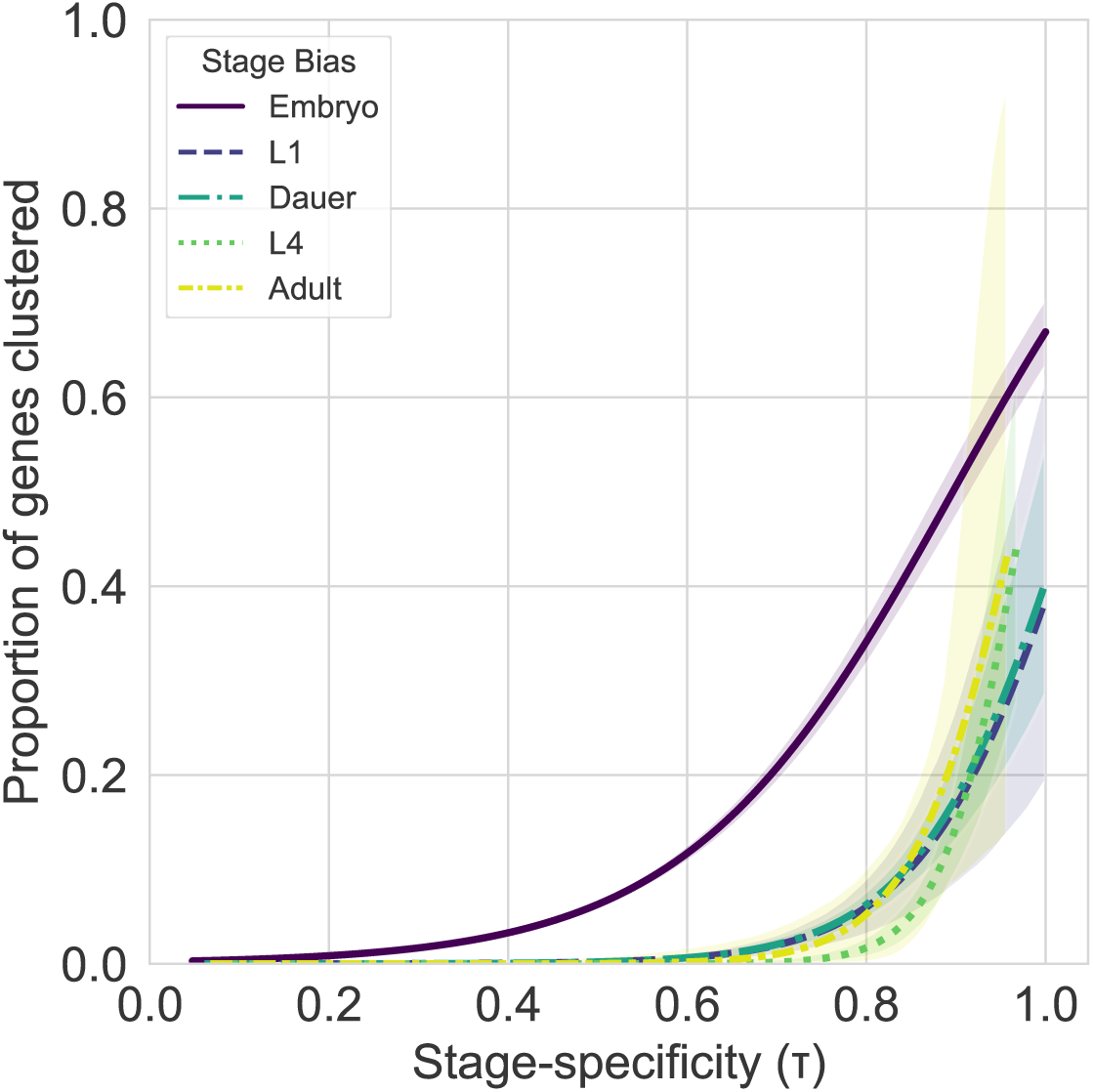
The proportion of stage-specific genes that are in stage-biased increases with stage-specificity and varies between life stages. Each line represents the logistic model that describes the proportion of genes that are in stage-biased regions (y-axis) as a function of stage-specificity (x-axis). Shaded regions around the lines represent the 95% confidence interval.

To mirror the analyses presented in the main text, we then examined the potential for gene duplication to contribute to non-random patterns of stage-biased regions. This showed that nearly 80% of genes within stage-biased regions did not have any detected homologs (Figure S5A). Furthermore, when using a larger window size we found that the non-homologous ratio was negatively correlated with gene density. This pattern was not apparent when using a small window size (*ρ* = 0.002, *p* = 0.98) (Figure S5B). Since using a larger window (100kb) 1) allowed for better quantification of gene duplications in stage-biased regions and 2) was more consistent with known dynamics of *C. elegans* genome biology, we focus on said analysis in the main text. Overall, using a smaller window seems to have broken down the larger stage-biased and duplicate-rich regions into smaller regions.

**Figure S6:**
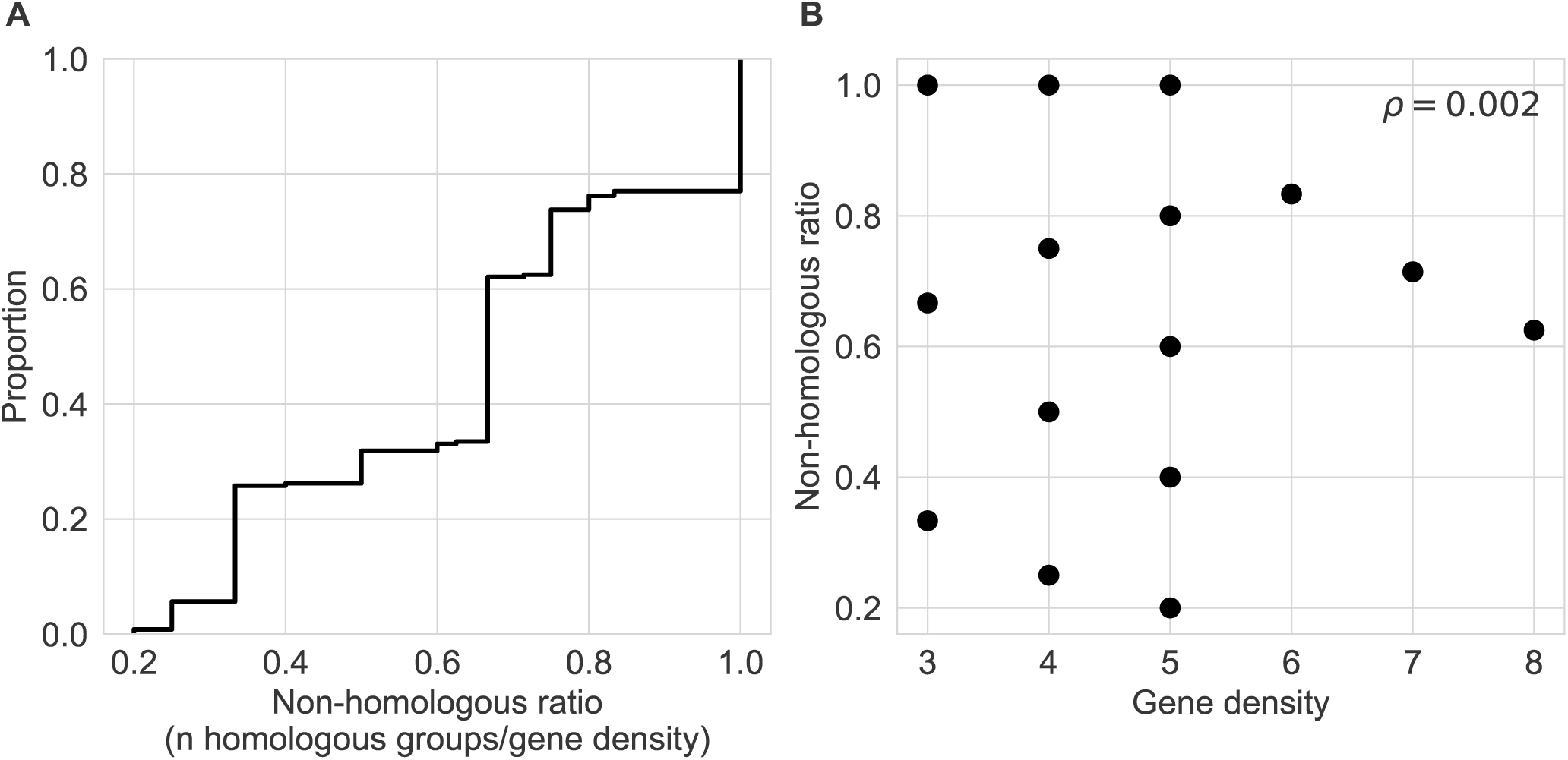
Patterns of expression similarity between genes in stage-biased regions can be partially but not entirely explained by gene duplication. A) An empirical cumulative density plot showing the distribution of non-homologous ratios across stage-biased regions. Non-homologous ratios represent the number of homologous groups represented in a stage-biased region relative to the number of genes. A non-homologous ratio of 1 indicates that all genes within a stage-biased region are non-homologous. B) Non-homologous ratios as a function of gene density for stage-biased regions.

#### 1.3 Logistic model summaries

**Table S1:** Logistic model summaries from the main text.

| Embryo |  |  |  |  |
| --- | --- | --- | --- | --- |
| Coefficient | Estimate | Standard Error | z value | p |
| Intercept | -4.99178 | 0.09606 | -51.97 | $< 2e - 16$ |
| $\tau$ | 5.22699 | 0.16005 | 32.66 | $< 2e - 16$ |
| L1 |  |  |  |  |
| Coefficient | Estimate | Standard Error | z value | p |
| Intercept | -10.693 | 1.271 | -8.414 | $< 2e - 16$ |
| $\tau$ | 11.360 | 1.616 | 7.030 | $2.07e - 12$ |
| Dauer |  |  |  |  |
| Coefficient | Estimate | Standard Error | z value | p |
| Intercept | -7.8019 | 0.6000 | -13.00 | $< 2e - 16$ |
| $\tau$ | 6.5640 | 0.7899 | 8.31 | $< 2e - 16$ |
| L4 |  |  |  |  |
| Coefficient | Estimate | Standard Error | z value | p |
| Intercept | -15.848 | 3.202 | -4.95 | $7.43e - 7$ |
| $\tau$ | 14.386 | 3.727 | 3.86 | $1.14e - 4$ |
| Adult |  |  |  |  |
| Coefficient | Estimate | Standard Error | z value | p |
| Intercept | -9.808 | 2.048 | -4.790 | $1.67e - 6$ |
| $\tau$ | 10.023 | 2.482 | 4.038 | $5.39e - 5$ |

**Table S2:** Logistic model summaries from the supplement.

| Embryo |  |  |  |  |
| --- | --- | --- | --- | --- |
| Coefficient | Estimate | Standard Error | z value | p |
| Intercept | -6.1059 | 1.1281 | -47.68 | $< 2e - 16$ |
| $\tau$ | 6.8110 | 0.1961 | 34.73 | $< 2e - 16$ |
| L1 |  |  |  |  |
| Coefficient | Estimate | Standard Error | z value | p |
| Intercept | -11.901 | 2.001 | -5.947 | $2.73e - 9$ |
| $\tau$ | 11.440 | 2.469 | 4.633 | $3.61e - 6$ |
| Dauer |  |  |  |  |
| Coefficient | Estimate | Standard Error | z value | p |
| Intercept | -11.966 | 1.083 | -11.05 | $< 2e - 16$ |
| $\tau$ | 11.579 | 1.332 | 8.69 | $< 2e - 16$ |
| L4 |  |  |  |  |
| Coefficient | Estimate | Standard Error | z value | p |
| Intercept | -22.291 | 3.151 | -7.074 | $1.50e - 12$ |
| $\tau$ | 22.796 | 3.578 | 6.371 | $1.88e - 10$ |
| Adult |  |  |  |  |
| Coefficient | Estimate | Standard Error | z value | p |
| Intercept | -16.227 | 4.878 | -3.327 | $8.78e - 4$ |
| $\tau$ | 16.669 | 5.637 | 2.957 | $3.104e - 3$ |

## Notes

https://github.com/gabe-dubose/celga

### Competing Interest Statement

The authors have declared no competing interest.

